# Towards XNA molecular biology: Bacterial cell display as a robust and versatile platform for the engineering of low affinity ligands and enzymes

**DOI:** 10.1101/2020.01.06.896050

**Authors:** Eszter Csibra, Marleen Renders, Vitor B. Pinheiro

**Affiliations:** University College London, Department of Structural and Molecular Biology, Gower Street, London WC1E 6BT, UK; Rega Institute for Medical Research, Herestraat, 49 box 1030, KU Leuven, 3000 Leuven, Belgium; Institute of Structural and Molecular Biology, Birkbeck College, University of London, Malet Street WC1E 7HX, UK

## Abstract

Although directed evolution has been remarkably successful at expanding the chemical and functional boundaries of biology, it is limited by the robustness and flexibility of available selection platforms – traditionally designed around a single desired function with limited scope for alternative applications. We report SNAP as a quantitative reporter for bacterial cell display, which enabled fast troubleshooting and systematic development of the selection platform. In addition, we demonstrate that even weak interactions between displayed proteins and nucleic acids can be harnessed towards specific labelling of bacterial cells, allowing functional characterisation of DNA binding proteins and enzymes. Together, this establishes bacterial display as a viable route towards the systematic engineering of all ligands and enzymes required for the development of XNA molecular biology.

## Introduction

Modern molecular biology has been built on nucleic acid processing enzymes isolated from nature – enzymes and binding proteins optimised for *in vivo* activity, well-adapted to natural nucleic acids and, in most cases, able to discriminate between RNA and DNA. Systematic mining of biological diversity, coupled to optimization of *in vitro* reaction conditions, has built a diverse range of ubiquitously used tools including nucleases, polymerases, ligases and kinases.

Detailed biochemical and structural information coupled to directed evolution strategies have enabled the enhancement of natural enzymes, modulating enzyme activity^[1]^ and even changing substrate specificity^[2,3]^. Driven by the requirements of biotechnological applications, DNA polymerases remain the most well-studied and most heavily engineered nucleic acid processing enzyme, with multiple viable directed evolution platforms available, covering *in vivo*, *in vitro*, *ex vivo* and *in silico* strategies^[4,5]^.

Polymerase engineering has also been central for the development of novel genetic materials based on synthetic or xenobiotic nucleic acids (XNAs). XNAs are nucleic acid analogues modified in at least one of the chemical moieties that define a nucleic acid: nucleobase, sugar and phosphate backbone^[6]^. Those modifications can alter the biophysical properties of the polymer, as well as its biological and chemical stability, making XNAs relevant for therapeutic aptamer developmen^[7–10]^, for nanotechnology^[11,12]^ as well as for xenobiology. XNAs that retain specific and unambiguous base-pairing potential are possible genetic materials, whether introduced alongside the natural system^[10,13–15]^ or as independent episomes^[16–18]^.

While simple *in vitro* XNA applications, such as aptamer selections, can be accessed with current DNA-dependent XNA polymerases (XNA synthases) and XNA-dependent DNA polymerases (XNA reverse transcriptases)^[2,9,19]^, more advanced XNA applications, whether *in vitro* or *in vivo*, require improved XNA polymerases as well as multiple other XNA-specific activities. In the particular case of *in vivo* XNA applications, the import of XNA precursors^[13,20]^, the assembly and maintenance of stable XNA episomes, and a viable catabolism for XNA by-products^[21]^ – all capable of function in the complex cellular environment and orthogonal to the cell (i.e. not interacting with the natural nucleic acids nor natural nucleic acid processing enzymes) – are the key challenges that need to be addressed. *In vivo* XNA platforms are expanding the chemical diversity of current biological molecules and have the potential to open new routes towards enhanced biocontainment, and to give novel insights into the boundaries of life^[17,22]^.

Systematic engineering of XNA molecular biology is possible, even in the absence of detailed biochemical and structural information, through directed evolution^[5]^: either via the development of multiple specialist selection platforms, each designed to target the individually desired functions, or via the development of robust platforms that can be easily adapted for the selection of multiple activities.

Here, we establish a flow cytometry-based reporter platform and use it to systematically develop bacterial cell display as a robust directed evolution platform. In addition, we demonstrate that cell display can be used for screening and engineering multiple protein functions, including low affinity ligands and enzymes, making cell display an ideal platform for the systematic development of XNA molecular biology tools.

## Results

In this work, our aim was to develop a bacterial cell display method for engineering XNA binders and processing enzymes by directed evolution. Although multiple platforms for the display of peptides and proteins on the surface of bacterial cells have been developed^[23–27]^, there are few reporter systems available to benchmark and compare the different display platforms^[28]^, and none that can be used for the systematic development of bacterial cell display as a platform for selections based on affinity and fluorescence.

### SNAP as an optimisation tool for display-based selection platforms

A suitable reporter protein for cell display optimisation should be easily expressed in *E. coli*, with its expression and activity readily quantifiable in bulk (by SDS-PAGE) as well as across populations (by flow cytometry). It should allow rapid discrimination of subcellular location – to distinguish between intracellular and displayed proteins – and it should provide a clear readout against an inactive phenotype – to enable the selection optimisation.

In this context, we focused on the display of SNAP, a small engineered protein derived from a human DNA repair protein, O_6_-alkylguanine-DNA alkyltransferase, that has been successfully used as reporter and as development tool of selection platforms^[29]^. SNAP catalyses the transfer of any chemical moiety attached to a benzyl-guanine (BG) to its catalytic cysteine^[30]^, allowing its own specific labelling. Expression of SNAP is well-tolerated in bacterial (and eukaryotic) hosts, and SNAP has been shown to tolerate N- and C-terminal fusions^[30]^. Crucial for its role in the development of bacterial cell display platforms is the commercial availability of BG-linked reagents including cell-permeable and cell-impermeable fluorophores (which allow precise localization of SNAP in and on the cell), and biotin (which can be used to demonstrate selection).

There is significant variation on the size (e.g. peptide vs large enzymes) and topology (N-tethered, C-tethered or N-and C-tethered) of the protein passenger between the bacterial display systems developed to date. Among those, two stand out: autotransporters, particularly EspP, which tolerate large passenger proteins and achieve high expression levels in *E. coli*^[31–34]^, and LppOmpA, which has been used successfully in both scFv antibody and enzyme display^[23,35]^, and is derived from a synthetic fusion between the Lpp signal sequence and a truncated OmpA.

Starting with the EspP-based display (Fig. 1a), we sought to demonstrate that SNAP could be expressed on the *E. coli* surface and to quantify its expression. Optimisation of the expression parameters allowed high levels of expressed protein (Fig. 1b) on the surface of DH10β *E. coli* cells (estimated at 1,000-10,000 molecules per cell or higher from densitometry measurements on protein bands compared to standards; data not shown).

**Figure 1.**
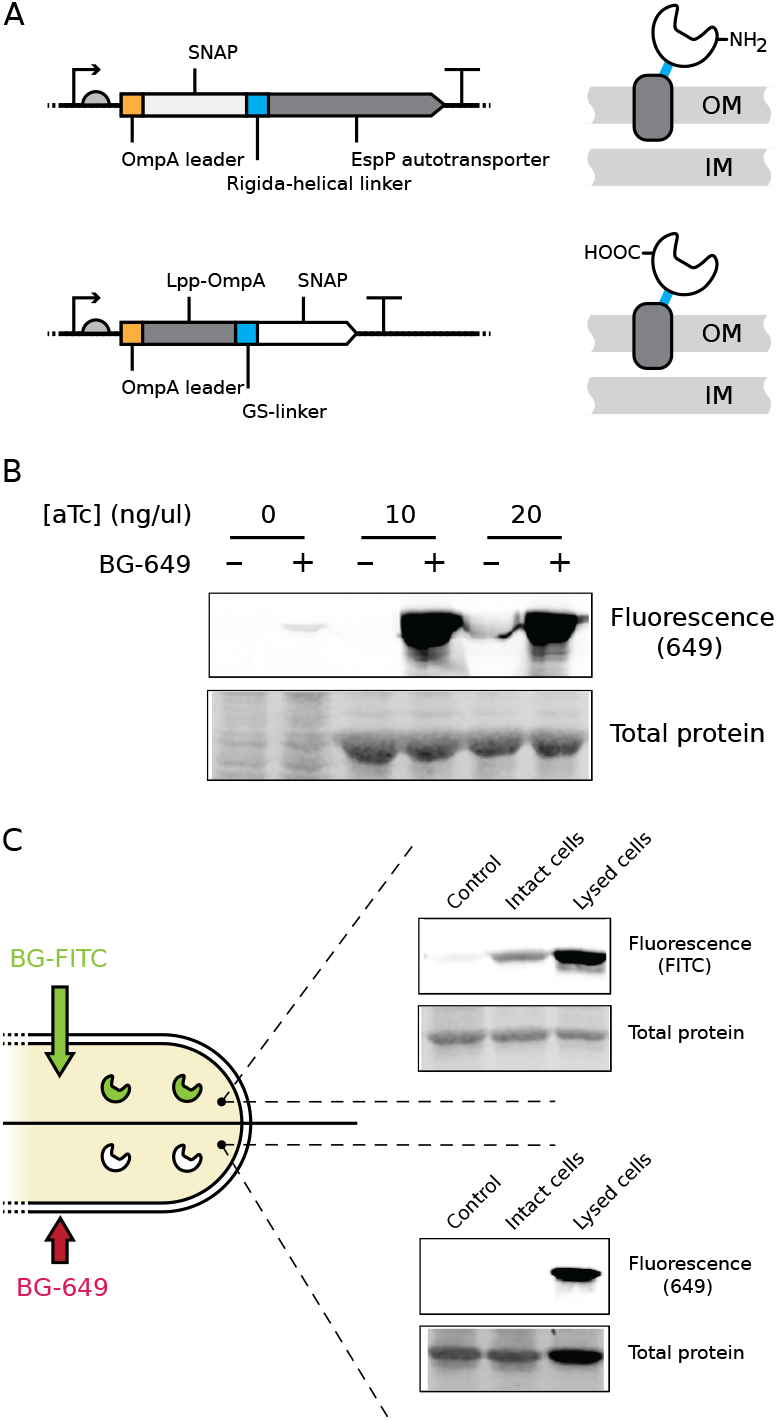
SNAP display. **A.** SNAP display variants. Cartoon of the SNAP-EspP (top) and LppOmpA-SNAP (bottom) expression constructs showing exposed N-terminal or C-terminal end of the protein of interest, respectively. **B.** SNAP expression on the cell surface monitored with membrane-impermeable benzylguanine-649 (BG-649), with and without anhydrotetracycline (aTc) induction and addition of the label. Insoluble protein fractions were separated by SDS-PAGE and imaged by fluorometry and post staining with Coomassie blue (Typhoon imager). **C.** Cytosolic SNAP expression (using pSNAP(T7-2)) monitored using membrane permeable BG-FITC labelling (top panel) and membrane impermeable BG-649 (bottom panel), showing mock labelled cells (Control), cells labelled while intact, and cells labelled post lysis. Soluble protein fractions were separated and imaged as above.

While protein localisation was typically validated by analysing the protein content of the insoluble fraction (SI Fig 1a-b), cell-impermeable SNAP substrates (Fig. 1b and c) enabled the accurate detection and quantification, by both SDS-PAGE and flow cytometry, of functionally active SNAP displayed on the surface of bacterial cells (Fig. 1b and SI Fig. 1c). Similar results were also obtained with LppOmpA fusions (Fig 1a, SI Fig. 1c).

Having demonstrated functional display of passengers, we focused on demonstrating the system was sufficiently robust for selection. Swapping the substrates used above for a BG-biotin conjugate allowed us to link the functional SNAP display to capture on streptavidin beads (Fig. 2a). Recovered cells were analysed for their genotype by diagnostic PCR (Fig. 2b) showing cell-density dependent enrichment of active over inactive clones of up to ~100-fold in one round: a demonstration that EspP systems can be adapted for selection of binding proteins.

**Figure 2.**
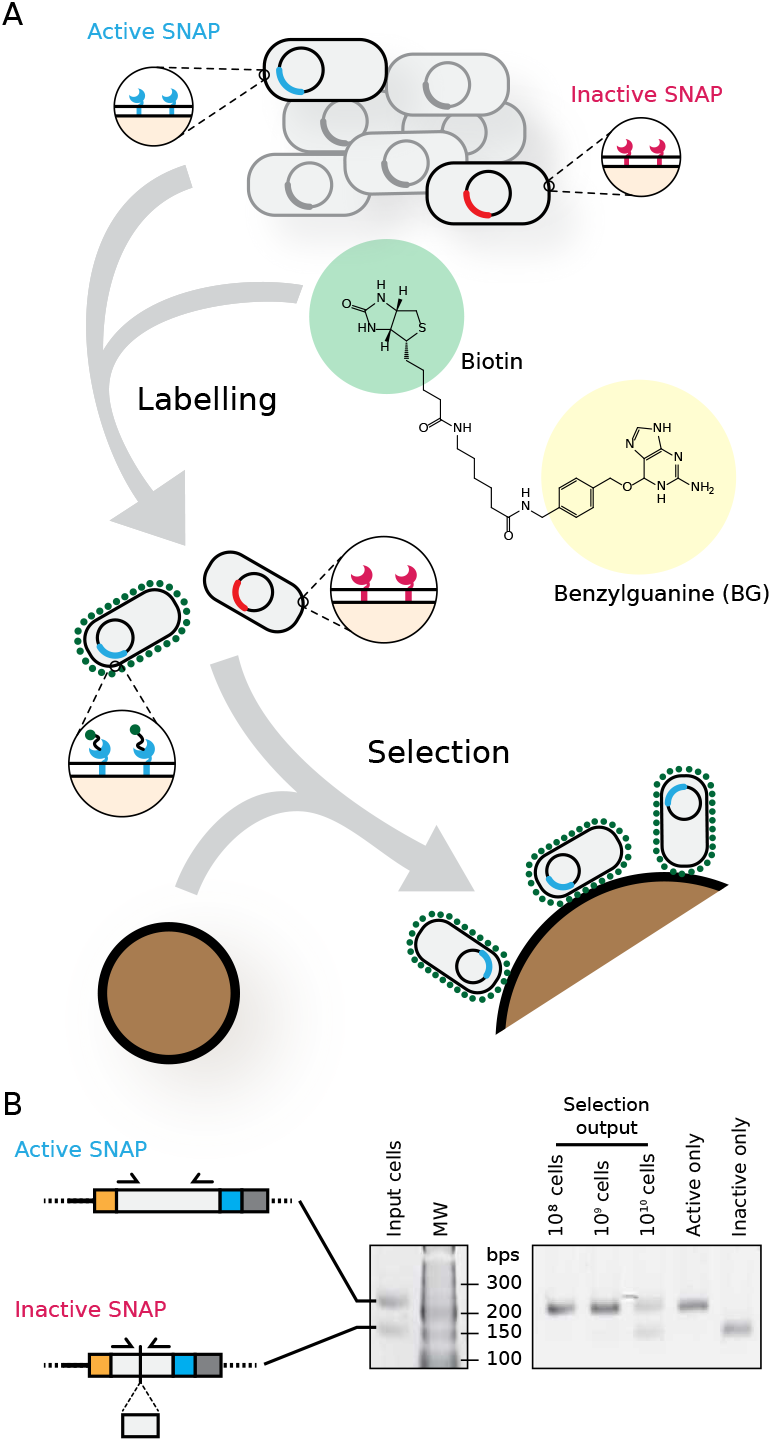
SNAP selection. **A.** Diagram of the SNAP selection method, using bead selection. Cell displaying Active and Inactive SNAP variants were induced, mixed, and labelled with BG-biotin. Streptavidin coated beads were incubated with these cells, unbound cells washed off, and bound cells analysed by differential PCR. **B.** Model selection PCRs. Inactive SNAP constructs were designed with a deletion (residues 83-100) that shortened the PCR product using the same set of primers. PCR products using plasmid DNA obtained from bead-bound cells (using range of input cell concentrations) are shown, next to Active or Inactive-only control PCRs, and the 1:1 input cell population.

The robust display of functional SNAP fusions and the efficient screening platforms were used to optimise expression conditions for EspP- and LppOmpA-linked fusions. They also enabled the development of protocols for the screening and selection of displayed passenger proteins, as a starting point for the engineering of DNA and XNA binding proteins.

### Cell display for the engineering of DNA/XNA binding proteins

Most *in vivo* replication systems rely on DNA binding proteins for correct function, whether by recruiting other proteins involved in replication, or simply inhibiting double strand formation^[36,37]^. In some systems, it can even be used to prime the replication itself^[38]^. Therefore, XNA binding proteins are essential for the development of a stable XNA episome both *in vivo* and *in vitro*. However, DNA binding proteins involved in DNA replication generally have weak affinity for their substrate, due to their requirement to bind DNA without sequence specificity, and functionally, many of them compensate for this by binding DNA co-operatively. Together, these are significant protein engineering challenges.

We focused on phi29 bacteriophage DNA binding proteins because of our ongoing effort to engineer phi29 XNA polymerases^[39]^ and because a simple DNA episome *in vitro* based on phi29 had already been established^[40]^. In addition to a DNA polymerase (p2) and a terminal protein (p3), two DNA binding proteins are known to be involved in episome maintenance and replication in phi29: p5 (single-stranded DNA binding protein) and p6 (double-stranded DNA binding protein)^[41]^. Both are known to bind DNA co-operatively and potentially as multimers^[42,43]^, effects that have been previously exploited in other selection platforms for the isolation of weak interactions^[44,45]^. Initial experiments, using different displayed DNA binding proteins, showed no interaction with DNA (data not shown), suggesting that proteins were not being correctly displayed or that the avidity effects due to high expression levels of displayed protein were not enough to stabilize the weak DNA-protein interactions.

As a result of this, we chose to focus on another phi29 nucleic acid binding protein, p16.7. Like p5 and p6, p16.7 is also a low affinity, co-operatively binding protein. Unlike p5 and p6, however, it binds both single and double stranded DNA and, it is itself a membrane-bound protein^[46–48]^. Specifically, its topology suggested that it would be amenable to display on the external face of the *E. coli* outer membrane and compatible with being tethered by its N-terminus^[49,50]^.

A truncated p16.7 (missing its transmembrane domain) was added as the C-terminal passenger to the LppOmpA-based display platform (Fig. 3a). Initial assays confirmed its high expression on the cell membrane (Fig. 3b) but initial binding assays again showed no signal above background (data not shown). Since the interaction of DNA with p16.7 is known to be weak, we considered if the expected high off-rate of binding could interfere with detection within the experimental time frame. We thus sought to stabilize any transient interaction between the DNA and the DNA binding proteins.

**Figure 3.**
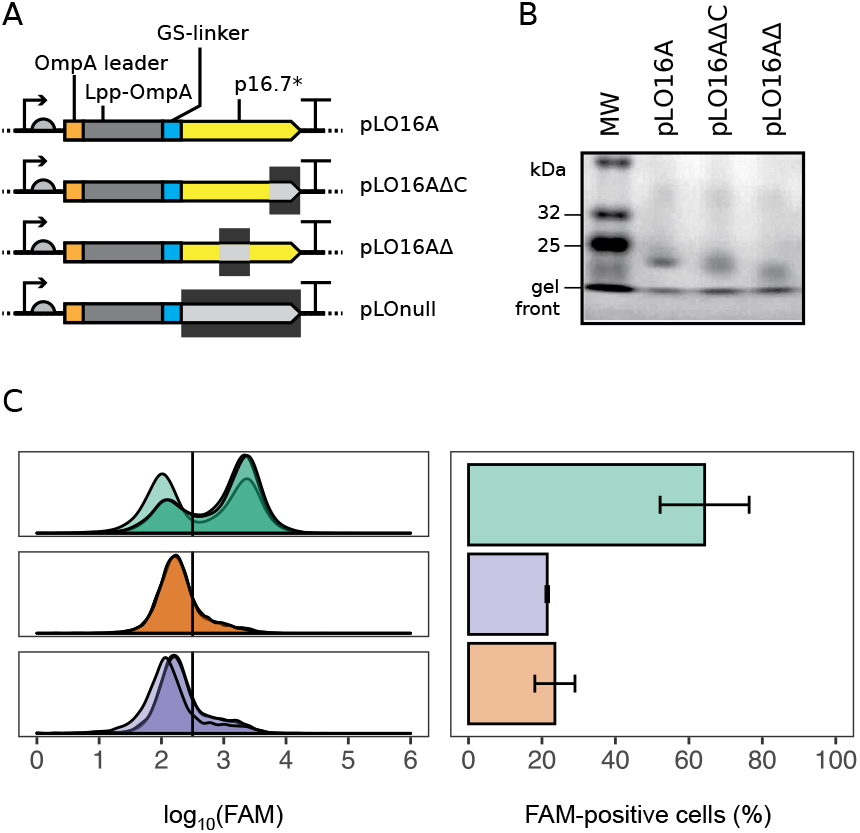
DNA binding protein display using p16.7. **A.** LppOmpA-linked p16.7 and its variants, showing leader sequence (orange), LppOmpA (grey), linker (blue), p16.7 variant (yellow) and deletion, if applicable (dark purple). The ‘wild type’ p16.7 variant (p16.7A) consisted of the full-length p16.7 sequence without the transmembrane domain (residues 22-130). Residues of the wild-type p16.7A remaining in the other variants were: LO16A_deltaC: 22-121; LO16A_delta: 22-94, 124-130; LO-null: none. **B.** Expression of LO16A variants, showing insoluble fractions separated by SDS-PAGE and stained with Coomassie blue. **C.** *Left*: DNA binding activity of LO16A variants. FAM-labelled DNA (EC98b_chol) was added to cells and incubated at 30°C (30 min, with shaking), before washing and analysis by flow cytometry (Attune NxT cytometer, BL1-H channel). The indicated 1D gate separates cells considered ‘negative’ vs. ‘positive’, assigned by adding a gate that put >95% control sample cells into the negative population. This gate acts as an indicator of the proportion of active and inactive cells that would be sorted by FACS in a selection experiment based on fluorescence rather than biotin/bead binding. Overlapping traces show data from triplicate samples. *Right*: Quantification of the proportion of cells with fluorescence above the gate threshold value. Error bars indicate standard deviation.

While multiple strategies are available to promote crosslinking between the bacterial cell and incoming DNA^[51–54]^, we opted for a commercially available cholesterol modification^[55,56]^. The modification alone is not sufficient to allow DNA binding to bacterial cells with intact membranes (SI Fig. 2a), but it enabled labelling of the bacterial cells in the presence of displayed DNA binding proteins (e.g. Fig. 3c). Labelling was fast, stable (within the experimental time frame), accessible to externally added DNase and could be outcompeted by unlabelled DNA (SI Fig 2b-e).

By using displayed DNA binding proteins to facilitate cell labelling, we were able to detect DNA binding on all tested phi29 proteins, including those required for *in vitro* episome maintenance: p3 (terminal protein), p5 and p6 (SI Fig. 3), suggesting that the platform is general and that it can detect weak interactions in nucleic acid binders. Discrimination between functional and inactive proteins was best for p16.7 (Fig. 3c), with a higher fraction of the expressing bacterial population labelled, potentially due to its evolutionary adaptations at binding DNA on a membrane surface. Displayed p16.7 also bound RNA (SI Fig 4).

The ability to display DNA binding proteins on the *E. coli* cell surface and detect even low affinity interactions with nucleic acids, while stably compartmentalising the genotype within the host cell, enables this platform to be used for the high-throughput screening of novel DNA and XNA binding proteins by flow cytometry. In fact, we found that our platform enables the detection and identification of low affinity interactions (such as phi29 proteins + DNA) that are hard to capture *in vitro*, due to the requirement to purify such proteins to challengingly high concentrations). It also opens up a route to attaching and stabilising nucleic acid substrates that might enable the engineering of DNA/XNA processing enzymes, as long as anchored nucleic acids can function as effective substrates for the displayed enzymes.

### Cell display for the engineering of DNA/XNA processing enzymes

By placing an enzyme on the outer surface of the bacterial cell, a key shortcoming of *in vivo* and certain *in vitro* selection platforms can be bypassed: displayed enzymes are not exposed to cellular metabolites and cellular proteins are not exposed to the reaction conditions of selection. Such reduced cross-reactivity is expected to increase the efficiency of enzyme selection.

For the demonstration that the systematic engineering of XNA molecular biology is possible on the cell surface, we focused on the display of phi29 DNA polymerase and T4 DNA ligase. Alongside polymerases, ligases are useful molecular biology tools and one of the essential functions needed to generate an XNA episome. DNA ligases have been reported to ligate some XNAs under forcing conditions^[57,58]^ but validated platforms for ligase engineering are limited^[59]^.

Preliminary assays showed that T4 DNA ligase (T4DL) was expressed at lower levels than the shorter DNA binding proteins from both display platforms. Nevertheless, increased levels could be achieved by expressing the T4DL fusions in C41(DE3) cells (hereafter ‘C41’), which was not solely due to its lack of the cell surface protease OmpT^[60]^ (SI Fig. 6B). A similar enhancement was observed when expressing the phi29 polymerase p2 (67 kDa, resulting in fusions of 102 kDa) on the cell surface (SI Fig. 5). Ligase display from EspP, although functional, was consistently poorer than LppOmpA-displayed ligase at catalysing the cholesterol-labelled DNA insertion into the membrane (SI Fig. 6c). DNA binding using LppOmpA ligase was strong in both DH10β and C41 strains, and could not enhanced by a *dsbA* strain that was previously reported to increase functional display of enzymes containing cysteine residues^[61]^ (SI Fig 6d). These observations highlight that the bacterial membrane is a complex environment and that standardization of the platform for different target proteins may be limited. In strains where expression was observed, displayed phi29 polymerase bound both single and double-stranded DNA, as expected (SI Fig. 5), indicating the C41 strain may have general potential for the display of large enzymes approaching 70 kDa in size.

Displayed ligases were active and able to ligate a splinted substrate in solution, in clear contrast to a non-functional catalytic site mutant (K159D)^[62]^ (Fig. 4b). Using the splinted substrate assay, quantifications of T4 DNA ligase activity could be determined compared to purified standards (SI Fig. 7). Adapting the splinted substrates to incorporate a cholesterol, compatible fluorophores or biotin, enabled separate detection of cell labelling (DNA1, ROX) and successful ligation (DNA2, FAM or biotin) directly on the cell surface, in little over 15 minutes (Fig. 4c and SI Fig. 8).

**Figure 4.**
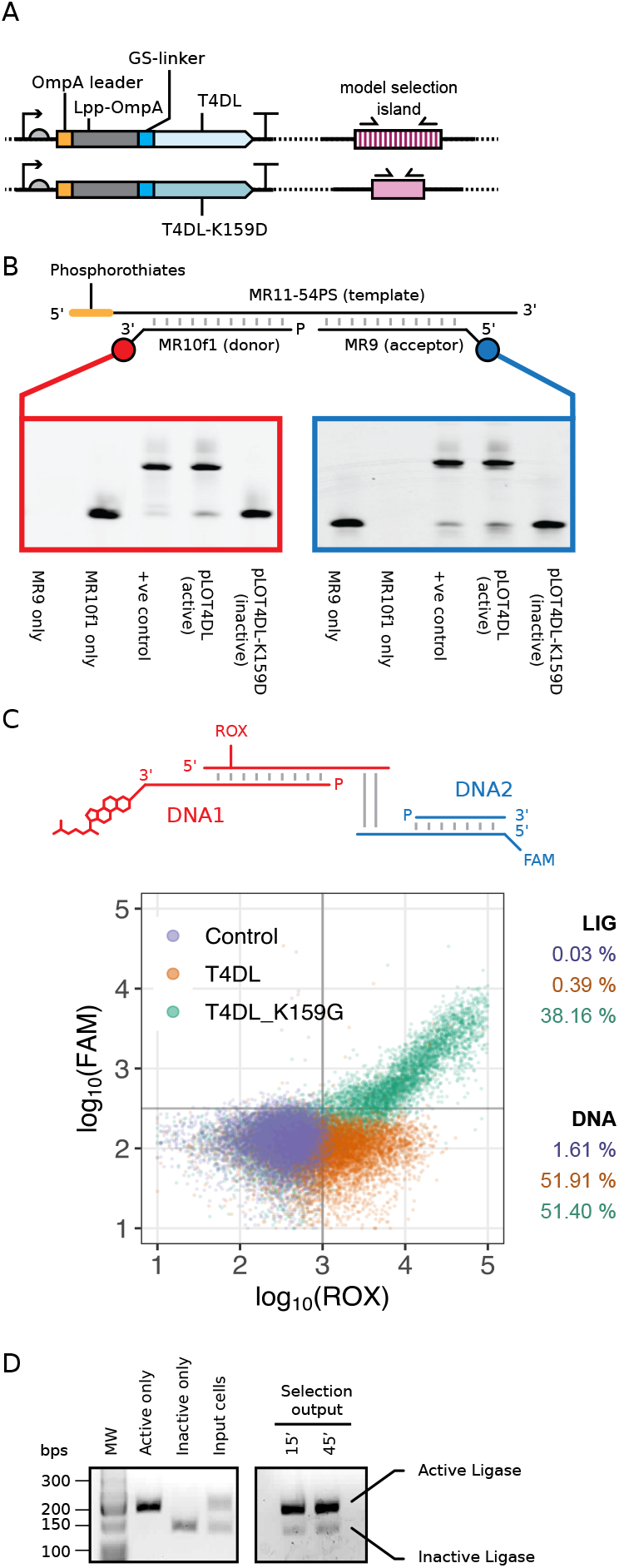
Ligase display. **A.** Ligase display constructs, showing leader sequence (orange), membrane protein (grey), linker (blue) and ligase variant (light blue). The wild type ligase (T4DL) was tested next to the active site mutant K159D. Active and inactive variants were differentiated by PCR by adding long (A) or short (I) model selection islands (pink, see Methods) downstream of the construct terminators to act as barcodes for PCR. **B.** Ligase function on the cell surface using soluble DNA substrates. The splinted ligation substrate, consisting of a splint (MR11-54PS), an acceptor strand (MR9, labelled with 5’ FAM) and a donor strand (MR10-f1, labelled with a 3’ Cy5), was mixed with cells displaying active or inactive ligase in ligase buffer. Following incubation at 37°C, cells were removed, and the DNA in the supernatant was separated using urea-PAGE and imaged by fluorimetry (Cy5, *left*, FAM, *right*). **C.** Ligase activity using cell surface-linked substrates. Ligation substrates consisted of DNA1, a duplex consisting of a ROX-labelled acceptor strand (top) and a donor strand (bottom) bound to cells via a cholesterol-linker, and DNA2, a similar duplex containing a FAM label (bottom strand). The overhangs were 4 nt (SI Table 2). Ligation activity was analysed by flow cytometry for active (T4DL) versus inactive (T4DL-K159D) cells. Plots show a sample of 1000 cells and indicate the proportion of cells considered positive for (i) DNA attachment (x axis, ROX label), and (ii) Ligation (y axis, FAM label). **D.** Model selection PCR of a ligase selection. Selections were carried out using a two-step DNA binding and ligation scheme as indicated above but using biotin (see SI Fig 7), and cells were mixed with streptavidin beads (as in Fig 2). Active and inactive variant cell populations isolated after stringent washing were differentiated by PCR as described in (a).

Model selection experiments, carried out under similar conditions, replacing FAM in the DNA2 primer with biotin to allow partition of the population by affinity (SI Fig 9), showed significant enrichment in one round of selection between active and inactive variants (Fig 4d), illustrating for the first time that bacterial cell display is a viable selection platform for the development of XNA-modification enzymes. Ongoing work is exploring applications of cell display not only for the development of DNA ligases for xenobiology as well as for other biotechnological applications.

## Discussion

### Systematic optimization of multiple parameters in method development and selection

The development of robust methods for the engineering of novel protein functions, such as needed for xenobiology, is reliant on optimisation tools that help researchers benchmark systems for user-specific goals. Previous work by Hollfelder and colleagues demonstrated that SNAP is a useful tool in troubleshooting selection methodology *in vitro*^[29]^, helping to optimise reaction conditions for each and every stage of selection. Here, we show that such an approach is also applicable to bacterial cell display. Efficient labelling of insolubly-expressed SNAP, using BG-linked fluorophores unable to diffuse into the bacterial cytosol, confirm not only that SNAP is functional but also that it is correctly displayed on the surface of the cells. Expression levels were high for both display platforms and the chemical labelling of SNAP was completed in minutes, leading to strong labelling of significant fractions of the bacterial population, which could be analysed by flow cytometry. Our approach establishes a fast and effective platform for the optimisation of expression conditions, including strains, induction times and inducer concentrations, all down to individual cell resolution. The commercial availability of different BG-linked fluorophores and BG-linked biotin also represent key advantages of the platform, enabling increased flexibility in multiplexed experiments (as the fluorophore to be conjugated to SNAP can be readily changed) and enabling a smooth transition from method development to selection, as shown above.

In addition, given the flexibility of labelling and of screening, bacterial cell display offers a unique opportunity to systematically optimise all steps in method development – a challenge regularly discussed in the field^[5,26,63]^ and not easily accessible to all selection platforms. Typically, the success of selections can be assessed by PCR enrichment in a model system (Fig. 2b and 4d), and while simple, this approach masks the complexity of parameters known to be involved in selection. We show that bacterial display is compatible with monitoring displayed protein and DNA levels (Fig 1b, 3c and 4c), and with introducing checkpoints [e.g. DNA binding separate from DNA ligation (SI Fig 8)] – allowing more complex selection strategies that take into account both enzyme and substrate levels with a cell-to-cell resolution.

### Stabilisation of low affinity interactions on the cell surface for the engineering of low affinity DNA binding proteins

While it is clear that selection can be used to isolate novel high affinity DNA binders^[64–66]^, low affinity interactions are not easy to engineer, despite being crucial in biology. Nonetheless, we demonstrate that weak interactions can be used to facilitate more stable ones, creating conditions amenable for selection.

Of the phi29 DNA binding proteins, p16.7 was the most successful (Fig. 3). Given its natural membrane-bound topology, it is likely that p16.7 could function unhindered on the *E. coli* outer membrane, enabling efficient labelling of the cells from a range of single-stranded and double-stranded, DNA and RNA substrates. Lower expression levels (which may derive from toxicity) and suboptimal tethering (e.g. protein fusions interfering with function) can account for the lower activity (both in % of labelled cells and in the strength of labelling of successful cells) seen with the other tested DNA binding proteins.

In addition to the engineering of DNA binding proteins, this platform may have other applications. For instance, displayed DNA binding proteins can be used to facilitate the incorporation of DNA nanostructures into biological membranes^[67]^, or to develop specific patterns of interactions between different bacterial populations^[55]^. Moreover, displayed proteins may provide an alternative biophysical route to quantitatively characterise weak protein:DNA or even protein:protein interactions, bypassing time consuming protein purification steps and technically challenging protein concentrations.

### Towards XNA molecular biology

Selection of nucleic acid processing enzymes with bacterial cell display has many of the constrains described above for DNA binding proteins – many of these enzymes bind nucleic acids weakly or non-specifically, and they must retain function while tethered to the cell surface against similarly tethered substrates. We show that both T4 DNA ligase and phi29 DNA polymerase, akin to the DNA binding proteins, can facilitate cholesterol-linked DNA attachment to the cellular membrane. The displayed DNA ligase remains active on DNA both in solution and when anchored to the membrane, underlining that despite the unnatural environment, both the folding of nucleic acid processing enzymes and their activity on DNA is robust on the *E. coli* surface. As with SNAP, labels can be readily swapped between fluorescence (for flow cytometry analysis) and affinity (for biotin-streptavidin selection), with flow cytometry again as a powerful tool to optimize all the steps in the method development, from validating target expression to selection stringency.

In all tested platforms and displayed proteins, high level expression invariably led to cell viability being compromised, with less than 1% cells remaining viable (data not shown). While cell viability is not necessary for *ex vivo* selections that do not rely on bacterial recovery, this would represent a significant challenge in the development of continuous evolution platforms^[68]^. In addition, it may limit the ability to build more advanced systems, such as those that display and select for the functions of multiple proteins at once.

### Conclusion

XNA molecular biology requires the systematic engineering of multiple new-to-nature activities and it relies on approaches capable of traversing those knowledge gaps, such as directed evolution. Selection platforms are the core of directed evolution and they differ significantly on throughput (number of variants that can concomitantly be sampled per round), efficiency (mean enrichment of maximum activity per round), flexibility (how easily a selection platform can be adapted for the selection of different catalytic activities), and on how isolated they are from other biological machinery.

Despite several potential advantages and successful demonstrations, bacterial cell display remains an underexploited selection platform. Here, we show SNAP display coupled to flow cytometry, provides a powerful tool for method development, allowing the optimization of throughput and efficiency. We demonstrate that DNA binding proteins and nucleic acid processing enzymes can be placed on the cell surface and their function can be monitored, despite their low affinity for a nucleic acid target, adding to the range of activities that can be selected in this platform.

Together, this demonstrates that the display platform is suitably flexible and sufficiently robust to be used in the systematic development of XNA molecular biology. Additionally, display protects the target protein from the cellular metabolites and enzymes that could interfere with selection, thus we anticipate that display will also prove key for the development of the mesophilic proteins required for the development of an XNA organism.

## Materials and Methods

### Bacterial strains, oligonucleotides and plasmids

Strains were obtained from NEB (DH10β), and as gifts from Renos Savva (Birkbeck, C41(DE3)), Valérie Pezo and Philippe Marlière (DH10β *ompT*), and Filipe Cabreiro (Imperial College London, K-12 BW25113 *dsbA* strain from the Keio collection^[69]^). Oligonucleotides were ordered from IDT. Plasmids were constructed as detailed below. All plasmids were cloned using Type IIS cloning or Gibson assembly (NEBuilder HiFi DNA assembly kit). Molecular biology enzymes and kits were obtained from NEB or Thermo Fisher Scientific, unless otherwise stated. SNAP reagents (BG-649, BG-fluorescein and BG-biotin) were obtained from NEB. Electrocompetent cells were used in all transformations.

### Constructs

Construct pNGAL97 was a gift from Prof. Arne Skerra (Technische Universität München, see ^[34]^). All cell display constructs were based on this vector. The SNAP tag sequence was obtained from NEB vector pSNAPtag(T7_2) (N9181S). The phi29 genes P3, P5 and P6 and expression constructs were gifts from Margarita Salas (Universidad Autónoma, Madrid). phi29 DNA polymerase p2 was derived from a synthesised N62D construct^[39]^. T4 ligase was a gift from Thermofisher Scientific.

For plasmids cloned using Type IIS method, the following protocol was followed. In each case, vector DNA (and insert DNA if applicable) were amplified by PCR using Q5 polymerase (NEB) and primers specified in Table 1, followed by DpnI treatment and purification (GeneJET PCR purification kit, Thermo Scientific). Amplified fragments were digested with AarI or BsaI (using all purified DNA from the previous step, with 3 µl enzyme in a 50 µl reaction volume, digested for 3 h at the recommended temperature), re-purified, ligated (T4 DNA ligase, 5 min at RT), and transformed into DH10β cells. For plasmids cloned using Gibson Assembly, insert fragments were commercially synthesised (gBlocks, IDT) and used directly (or amplified first by PCR, DpnI-treated and purified as above) with PCR-amplified vectors. Transformants were verified by sequencing with primer EC42. (Unless otherwise specified, protocols followed the manufacturer’s recommendations.) A list of oligonucleotides used is included in the supplementary information.

### Agarose-lithium acetate gels

Agarose gels (made in 10 mM lithium acetate) were run in 10 mM lithium acetate (agarose-LiOAc)^[70]^. Gels were run at 260 V (10 V/cm) for 10-20 min and visualised by SYBR Safe staining (Thermo Fisher).

### DNA annealing

DNA substrates for binding or ligation were pre-annealed by heat-cooling (95°C for 2 min, −0.1°C/s for 20 min to 20°C) in a thermocycler, or freeze-thawed (−20°C for 2 h or more, then thawed slowly on bench for 30 min), with 10 mM Tris•Cl (pH 7.5), 5 mM MgCl_2_.

### Expression of cell surface display constructs

A number of different strains and conditions were used for expression of cell display constructs, see text for details. A typical expression was as follows. Overnight cultures were diluted in 2TY-amp (100 µg/ml ampicillin, 1:100 dilution) and grown at 37°C with shaking (250 rpm) for about 2 h. Constructs were induced with 10 ng/ml anhydrotetracycline (aTc) and grown for a further 30 min to 2 h at 30°C (250 rpm). Cell growth was assessed by OD600 readings (absorbance at 600 nm), cells were harvested with washing in suitable buffer, and resuspended to a concentration of 10 OD/ml (approximately 1 × 10^10^ cells/ml). An aliquot of washed cells was frozen for SDS-PAGE analysis and the rest was stored on ice until use.

### Assessment of membrane protein content

Aliquots of cells were sonicated using a QSonica 700 sonicator at 100% amplitude for 2 min (30s on/30s off) at 4°C and insoluble fractions were collected by centrifugation (16,000x*g*, 30 min, 4°C). Pellets were taken up in 1x Laemmli’s buffer (62.5 mM Tris•Cl pH 6.8, 2% SDS, 20% glycerol, 100 mM dithiothreitol, 0.5 mM bromophenol blue) and separated by 8-15% SDS-PAGE (0.05 OD equivalent/lane) and stained with Instant Blue Coomassie Protein Stain (Expedeon).

### SNAP assays

Cells (in PBS: 137 mM NaCl, 2.7 mM KCl, 8 mM Na_2_HPO_4_, and 2 mM KH_2_PO_4_) were incubated with BG-649, BG-fluorescein or BG-biotin conjugates (known as “SNAP-Surface® 649”, “SNAP-Cell® Fluorescein” and “SNAP-Biotin” respectively, NEB, at a 1:500 dilution, 2 µM final concentration) for 5-15 min at RT or as indicated. Labelling was carried out pre-sonication for BG-649 and post-sonication for BG-fluorescein, and assessed by SDS-PAGE scanned using a GE Typhoon FLA 9500 at the appropriate wavelengths.

### Model selections with SNAP

SNAP-EspP2 and inactive (harbouring a 54-nucleotide deletion from the middle of the SNAP ORF introduced by primers EC38fwd/rev.) variants were expressed separately in DH10β strain, mixed at a 1:1 ratio, and subjected to a SNAP labelling assay with BG-biotin. Labelled cells were added to streptavidin beads (5 µl, Dynabeads, MyOne C1, Thermo Fisher) in binding buffer (TN-DBT, 50 mM Tris•Cl pH 7.5, 10 mM NaCl, 1 mM DTT, 0.1% BSA, 0.01% Tween 20) and put through selection in a KingFisher™ Duo Purification System. The protocol included a binding step (30 min, 37°C, medium shaking, collect beads 3x 5 s), 6 wash steps (5 min, medium shaking, collect beads 3x 1 s), and an elution step (into 50 µl PBS). Eluates (1 µl) were used as templates for diagnostic PCR analyses.

### Model selection PCR

Model selection PCRs were designed such that the product of one PCR (two primers) would be diagnostic to the activity of the underlying construct (Fig 2b). PCR reactions were conducted using MyTaq (Bioline) DNA polymerase, with the following cycling conditions: 95°C for 5 min, followed by 30 cycles of 95°C for 30 s, 57°C for 10 s, 72°C for 10 s, and a final incubation at 72°C for 5 min. Products were analysed on 2% agarose-LiOAc gels.

The diagnostic PCR targeted the difference in size between the active (complete) and inactive (deletion mutant) variants of SNAP. For subsequent constructs, a model selection island (MSI) was constructed downstream of the terminator in the vector of interest (Fig 4a). For each active and inactive variant, an Active or Inactive MSI was built from segments of active and inactive SNAP genes between the primers EC19 and EC20. These islands were bordered by translation insulator elements.

### DNA binding assayed using flow cytometry

Cells (50 µl at 10 OD/ml, typically in PBS) were incubated with relevant DNA (or RNA, 500 nM) substrate as specified in the text. Typically, incubations were carried out at 30°C, for 30 min with shaking at 700 rpm (Thriller, Peqlab). Cells were pelleted (4000x*g*, 5 min, 4°C), resuspended in wash buffer (TMd buffer with 10% BSA and 100 µg/ml yeast tRNA) and incubated for 5-10 min at 30°C. After washing, cells were resuspended in PBS and diluted 1:1000 in PBS for analysis by flow cytometry (Attune NxT, Thermo Fisher).

### Flow cytometry data analysis

Inline thresholding, cell gating and preliminary data analysis was handled using the Attune NxT software. Data from 10,000 cells were collected by manual gating of singlets during data collection, but for visual clarity, random samples of the data were used for dot plotting. Further data analysis was carried out using R. Singlets were selected from bulk data using the autoGate package^[71]^, and resultant data was plotted using R. Fluorescence signals from the following channels were analysed: in the BL1-H channel for FAM/FITC (ex. laser: 488nm, em. BP filter 530/30), YL2-H channel for ROX (ex. laser: 561 nm, em. BP filter 630/15), RL1-H channel for Cy5 (ex. laser: 638 nm, em. BP filter 670/14). Flow cytometry data was used primarily for method development but also with a view of using FACS sorting for selection. As such, the key measurement in many of the experiments was the proportion of cells scoring as above a 1D or 2D gate, where the gates represented a threshold value not occupied by the negative population and usable as a sorting gate during FACS. The gates were assigned in a position that binned >95% control sample cells into the negative population and acted as an indicator of the proportion cells that would be sorted by FACS in a selection experiment based on fluorescence rather than biotin/bead binding.

### *Ex vivo* ligase assays for bulk analysis

Cells were washed and resuspended in TMd (10 mM Tris•Cl pH 7.5, 10 mM MgCl_2_, 1 mM DTT) at 10 OD/ml. Aliquots (50 µl) were washed again and resuspended in T4 ligase buffer (NEB) containing pre-annealed nicked DNA substrate (MR9/MR10-f1/MR11-54PS or as specified, 500 nM). Ligations were carried out at 37°C for 30 minutes or as specified in the text. DNA was isolated from the supernatant fraction and 1 µl (equivalent to 0.5 pmol DNA) was loaded onto gels for analysis. Commercial (NEB) T4 DNA ligase (400U per reaction) was used in positive control reactions.

### *Ex vivo* ligase reactions using cell-immobilised substrates

Cells were washed and resuspended in TMd at 10 OD/ml. Aliquots (50 µl) were washed again and resuspended in TMd containing pre-annealed DNA1 (500 nM, with/without 10% PEG-8000). DNA binding (DNA1, 0.5 µM) was carried out at 30°C for 30 minutes, after which cells were washed twice (5 min, RT, 700 rpm) in 100 µl TM-block (TMd with 10% BSA and 100 µg/ml tRNA). DNA ligation (of DNA2 to DNA1) was carried out by resuspending washed cells in ligase mixture, containing DNA2 (1.5 µM) in buffer (standard T4 buffer supplemented with 10% PEG 8000 and 0.5 µM ROX-containing DNA) in a volume of 25 µl. Ligations were incubated at 37°C without shaking for 15 min to 1 h and washed again twice in TMd. Control reactions (where applicable) used equivalent DNA mixtures in equivalent buffer and final volumes, adding 1 µl T4 DNA ligase (NEB, M0202S). Cells were diluted in PBS for flow cytometry analysis.

## Acknowledgements

E.C. and V.B.P acknowledge support by the European Research Council [ERC-2013-StG project 336936 (HNAepisome)] and BBSRC [grant BB/N01023X/1 – (invivoXNA)]. The authors would like to thank Alex Fedorec for his assistance with R scripting for flow cytometry data analysis.

