## Supplementary figures and images for "Towards XNA molecular biology: Bacterial cell display as a robust and versatile platform for the engineering of low affinity ligands and enzymes"

### SI_figures

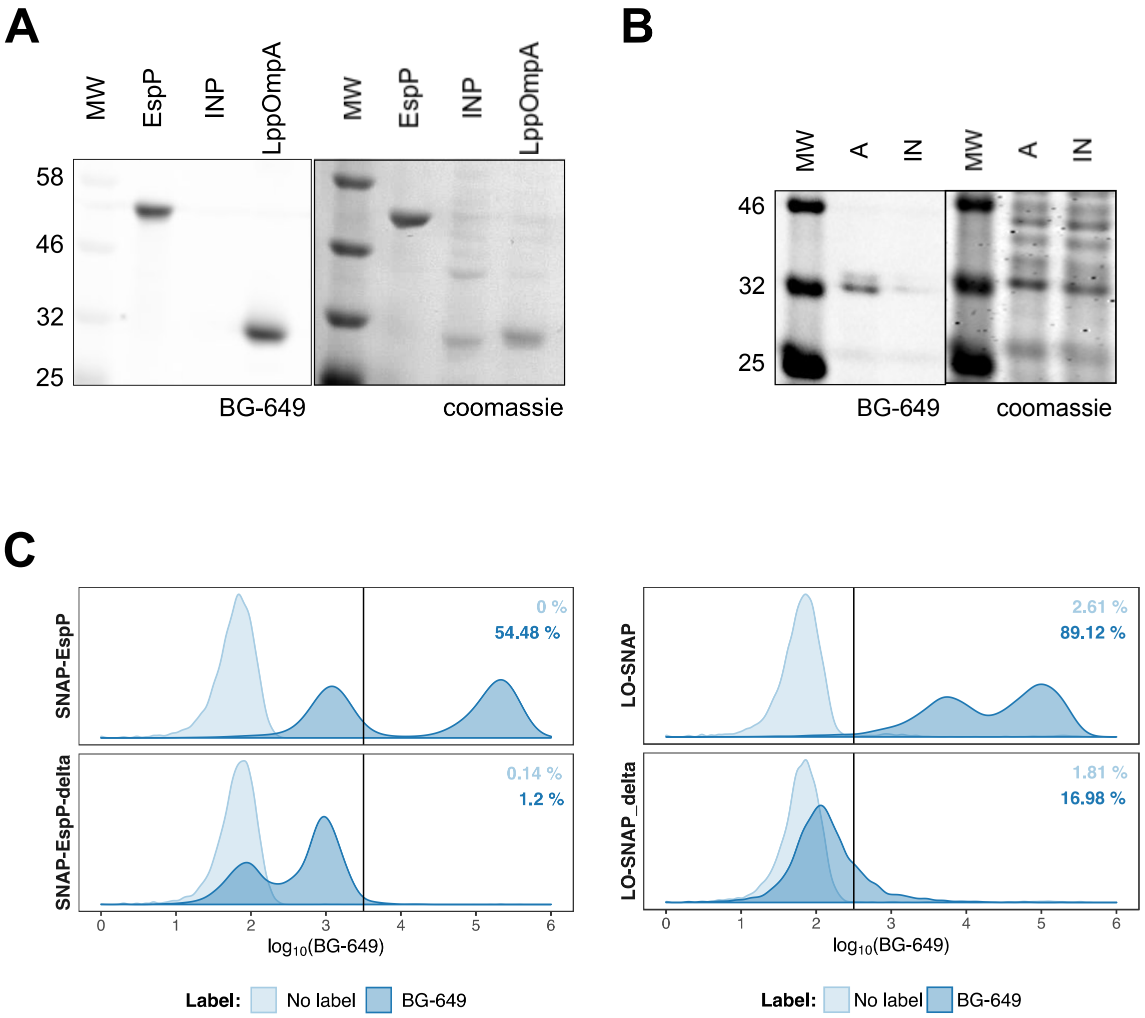

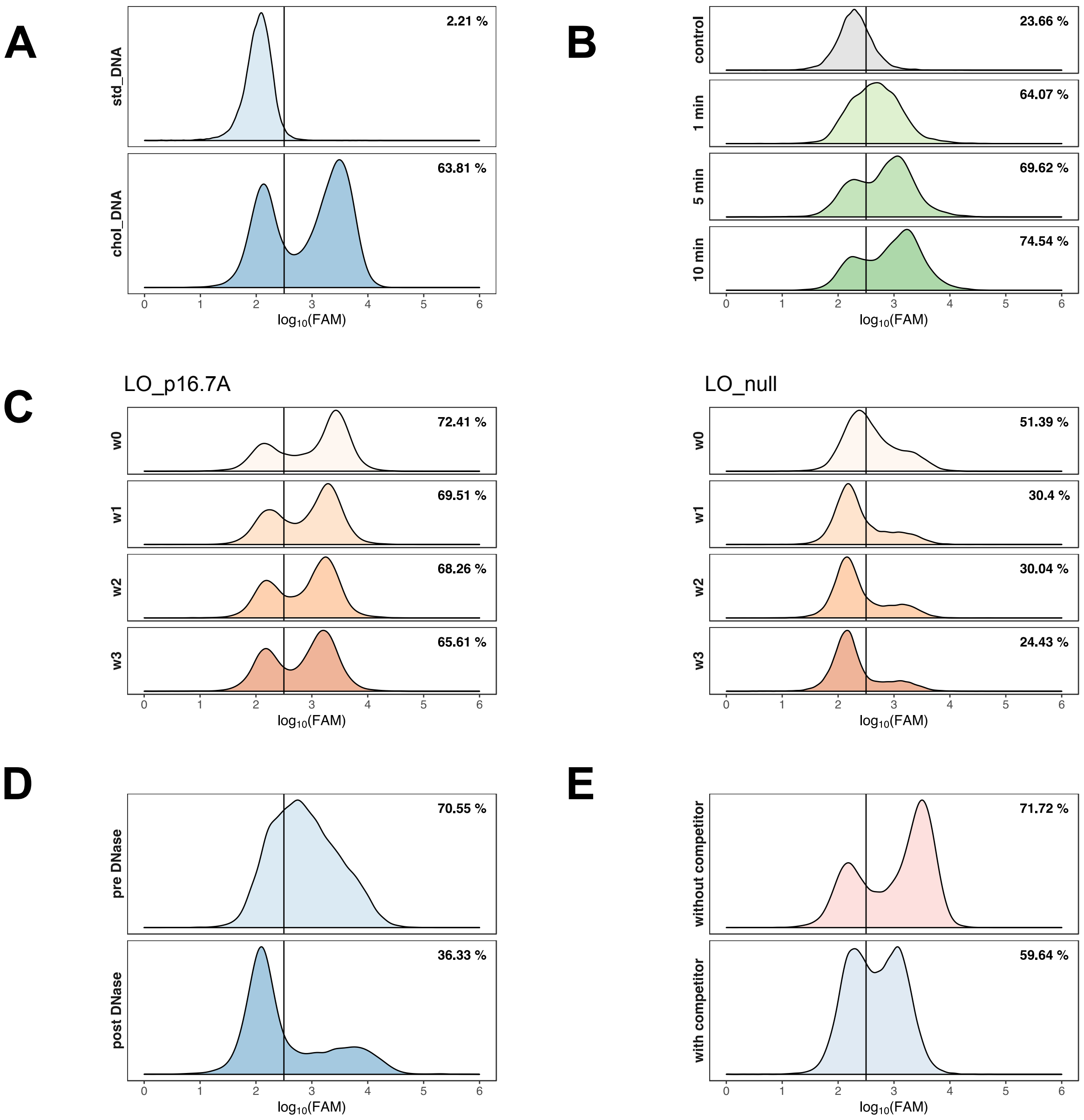

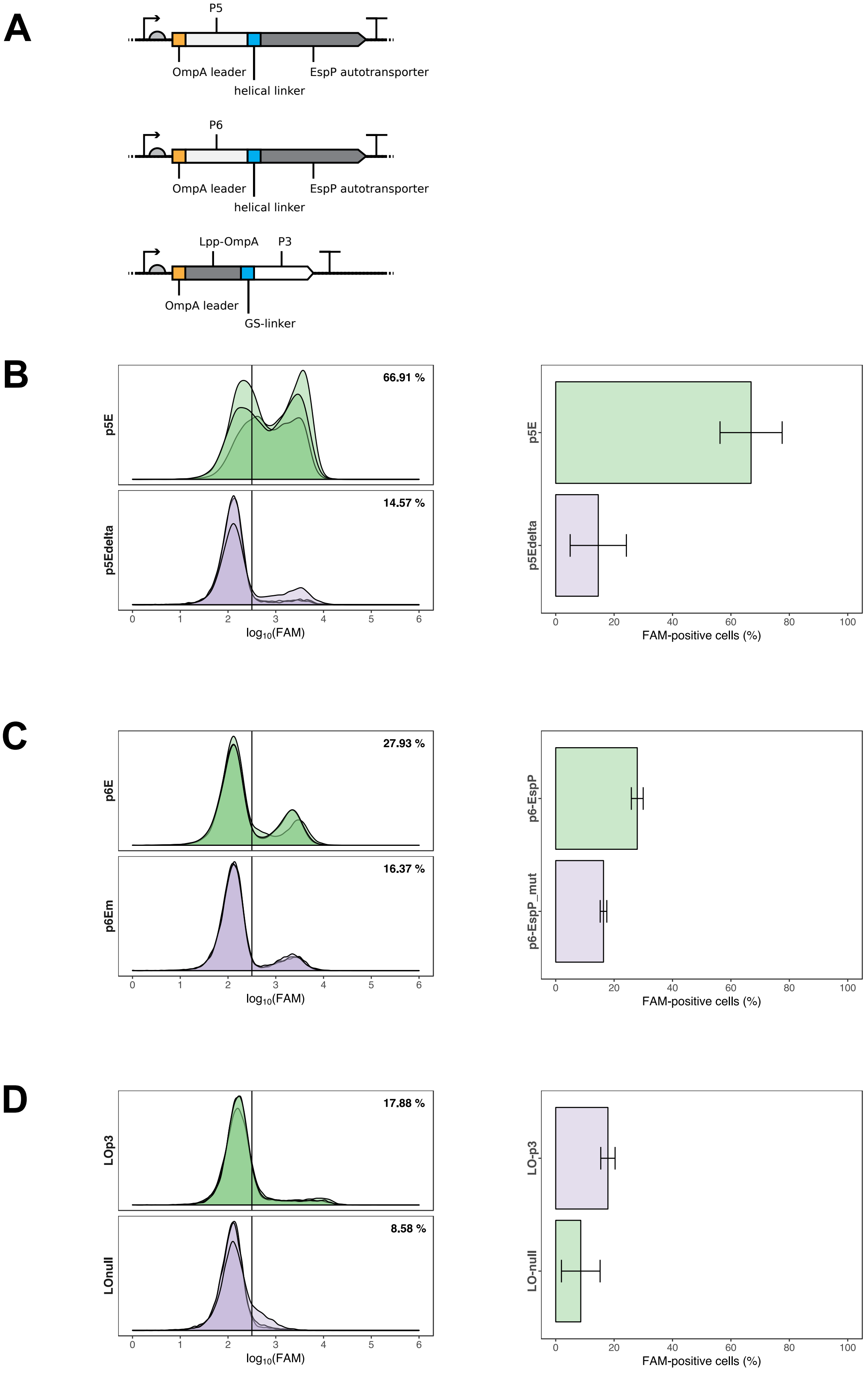

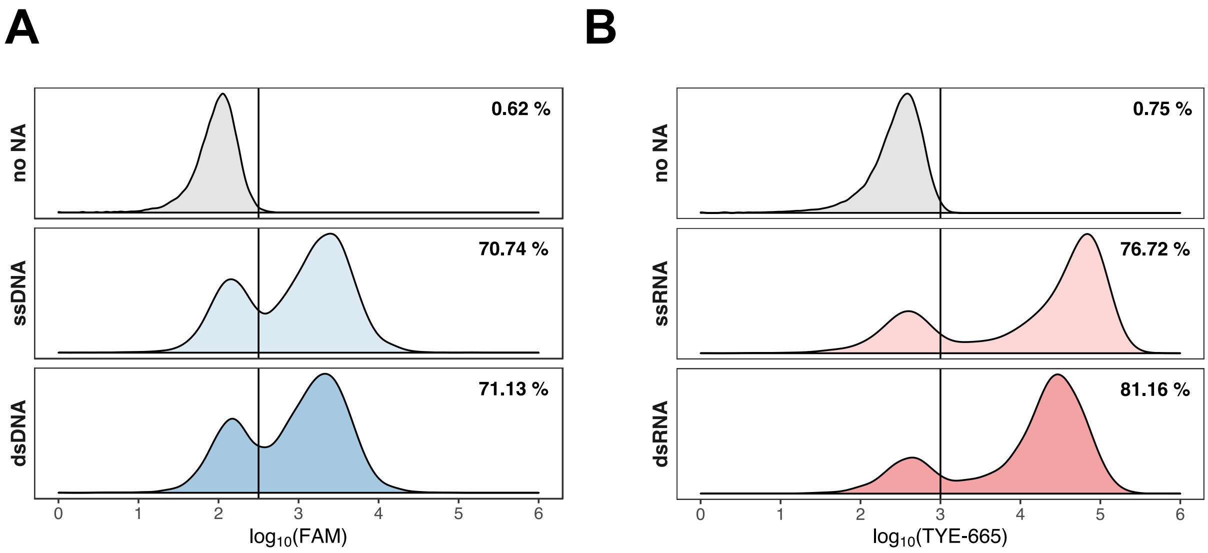

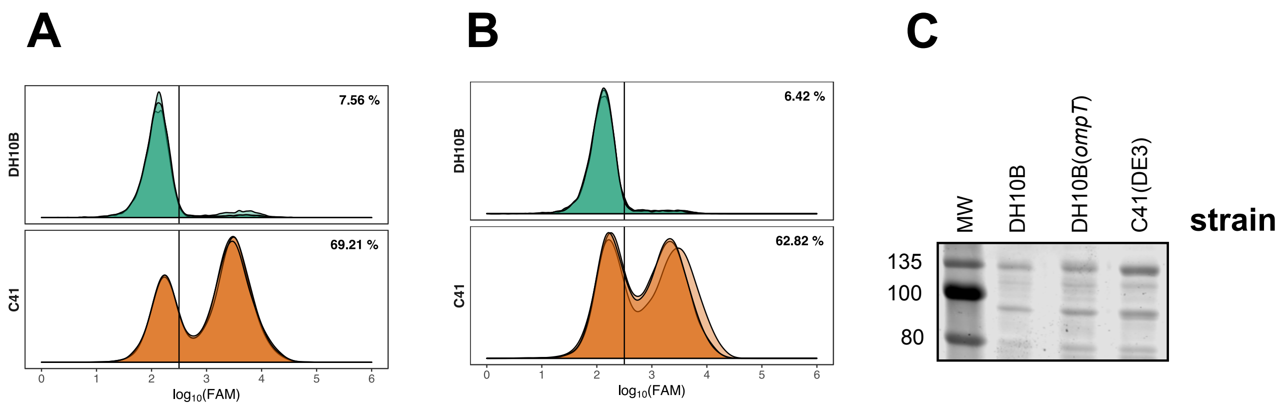

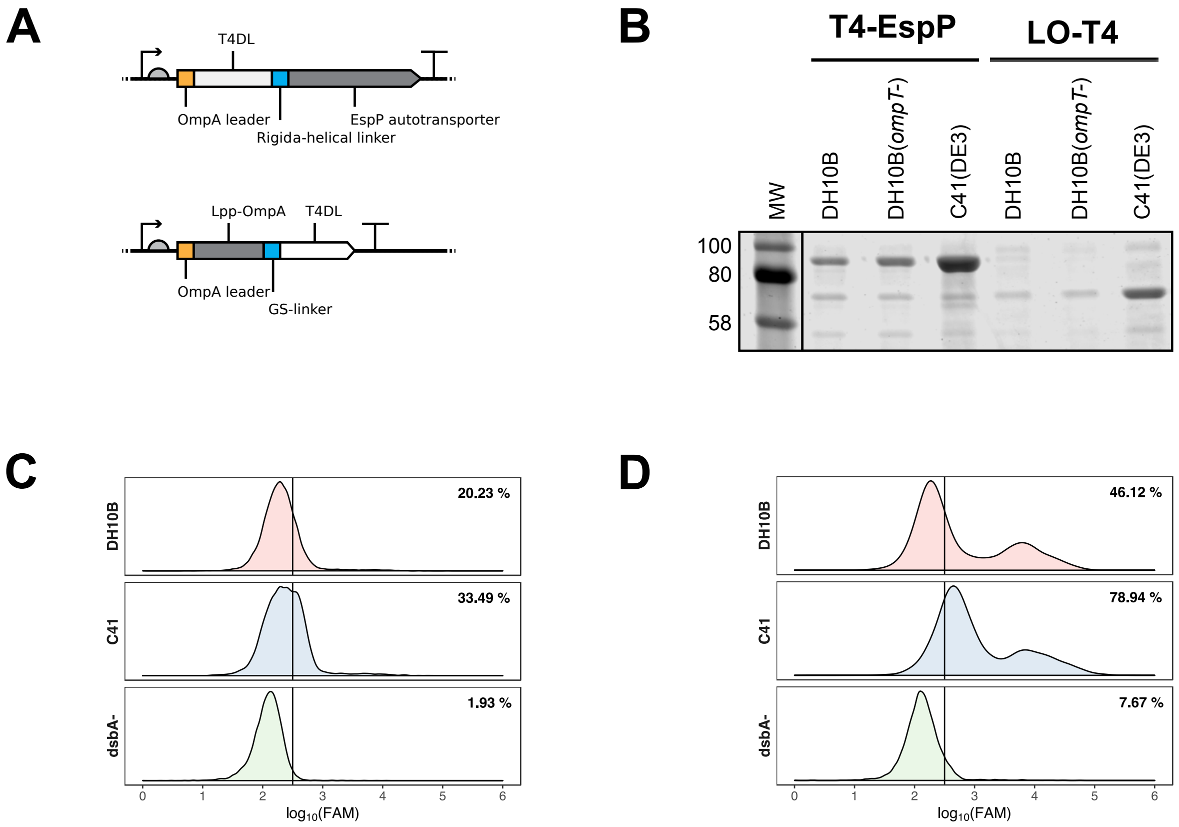

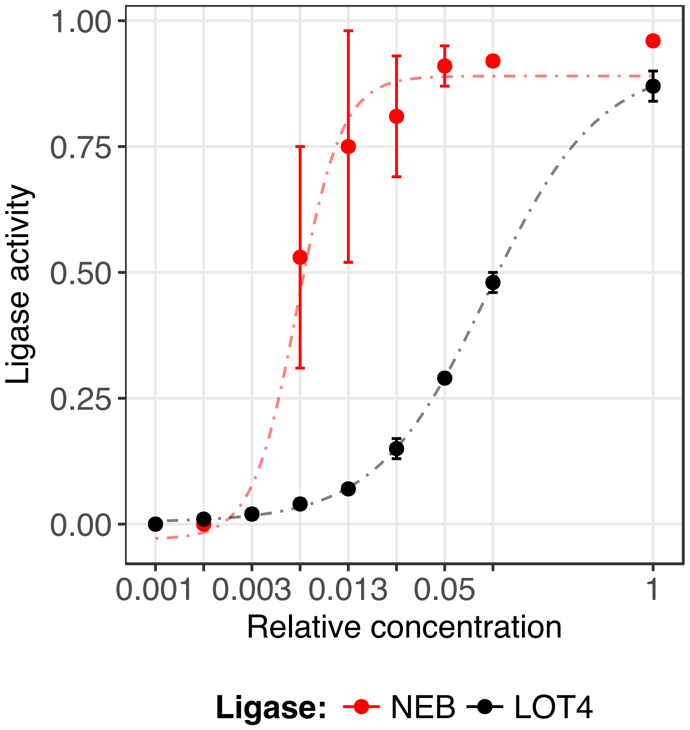

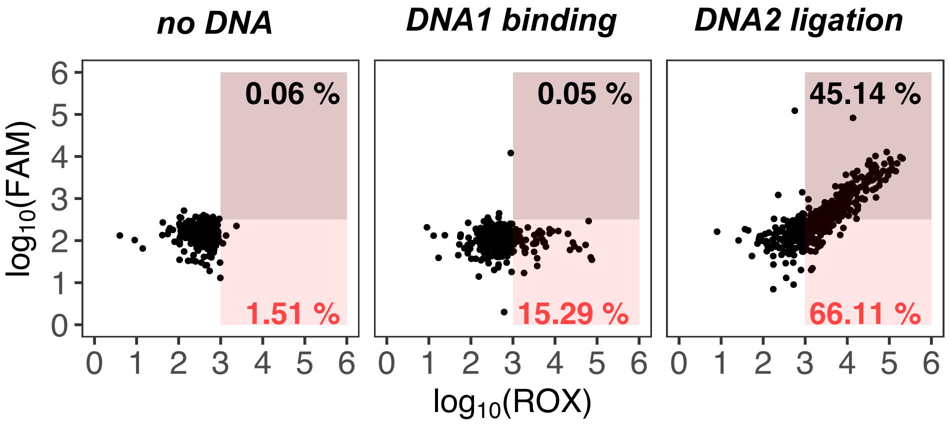

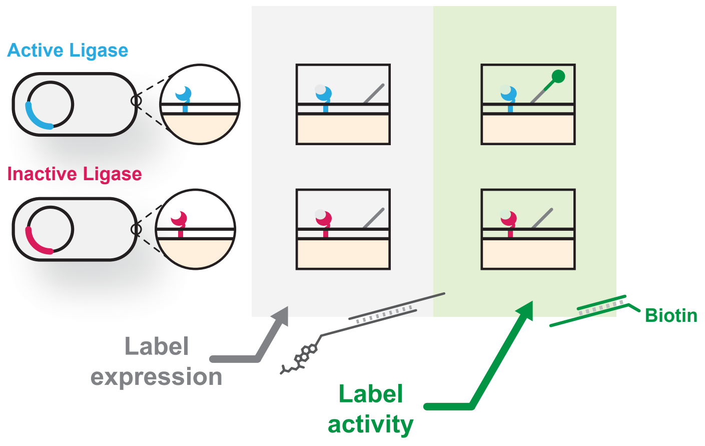
