## Supplementary material for "Towards XNA molecular biology: Bacterial cell display as a robust and versatile platform for the engineering of low affinity ligands and enzymes": SI_figure legends

Supplementary Figure Legends

**SF1. Cell surface SNAP expression and activity analysed by SDS-PAGE and flow cytometry. A.** SNAP expression (Coomassie panel) and activity (BG-649 panel) by display construct, stained as in Fig1b, showing insoluble protein fractions separated by SDS-PAGE and imaged by fluorimetry (Typhoon imager). Next to the EspP and LppOmpA systems, the ice nucleation protein (INP) system was also compared here. INP constructs structure resembled LppOmpA constructs in that the protein of interest is displayed on the C-terminal end of the fusion. **B.** SNAP expression (Coomassie panel) and activity (BG-649 panel) in active (SNAP-EspP) and inactive (SNAP-delta-EspP, missing residues 83-100) variants. **C.** SNAP activity as analysed by flow cytometry. Cells were incubated with BG-649, washed in PBS and analysed using an Attune NxT cytometer, RL1-H channel (10,000 cells/sample). Histograms show cell signal in the RL1 channel, while annotations (%) indicate the proportion of cells considered positive (that score over the indicated gate threshold value) for BG-649 binding. The gate acts as an indicator of the proportion cells that would be sorted by FACS in a selection experiment based on fluorescence rather than biotin/bead binding.

**SF2. Characterisation of cholesterol-linked DNA binding to cells with displayed p16.7 protein. A.** Comparison of FAM signal detectable on cells (pLO16A) mixed with cholesterol-labelled DNA (dsDNA: EC98a/EC98b_chol) versus unlabelled DNA (EC98a/EC98b). **B.** Timecourse of DNA binding to cells. Cells (pLO16A) were mock labelled (control) or incubated with DNA (EC98b_chol) for the indicated time (30^o^C, with shaking), before being washed in PBS and analysed. **C.** Analysis of the stability of cholesterol-DNA bound to cells. Cells (pLO16A, *left*; pLO16A_deltaC, *right*) were incubated with DNA (EC98b_chol, 30 min at 30^o^C with shaking), before being diluted for analysis (w0) or washed in PBS, recovered by pelleting and then diluted (w1-3) the indicated number of times. **D.** Analysis of DNA accessibility to the surrounding media. Cells (pLO16A) were incubated with DNA (EC98b_chol, 30 min at 30^o^C with shaking), washed, and subjected to DNase treatment (Turbo DNase, 37^o^C, 30 min), before washing and dilution for analysis. **E.** Analysis of the process of DNA:cell binding. Cells (pLO16A) were incubated with DNA (EC98b_chol, 30 min at 30^o^C with shaking), without (*top*) or with (*bottom*) the addition of excess competitor DNA (EC98b, without the cholesterol moiety) to ascertain whether the binding was driven by DNA:DNAbp interactions or cholesterol:membrane interactions. (The decrease in FAM signal suggests that the competitor, despite not having a cholesterol moiety, interferes with cholestol-DNA:cell binding, suggesting that binding of cholesterol-DNA is DNA:DNAbp driven).

Histograms show cell signal in the BL1-H channel, while annotations (%) indicate the proportion of cells considered positive (that score over the indicated gate threshold value) for DNA binding. Overlapping traces show data from triplicate samples; samples shown in singlet form are representative of multiple experiments.

**SF3. Cell surface display of DNA binding proteins from phi29. A.** Constructs used for DNA-binding protein display, showing leader sequence (orange), membrane protein (grey), linker (blue), DNA-binding protein (yellow). **B-D.** Assessmengt of DNA-binding by cells, using active and inactive variants of each protein, showing signal from FAM-labelled DNA on cells by flow cytometry (*left*) and the proportion of cells scoring as FAM-positive (*right*). **B.** p5 (ssDNA binding protein) display using EspP, with p5-delta (missing residues 44-61) as the inactive control. **C.** p6 (dsDNA binding protein) display using EspP, with p6-mutant (K3A, R7A, K11A, T12A) as the inactive control. **D.** p3 (ss/dsDNA binding protein) display using LppOmpA, with LO-null as the inactive control.

Cells were induced for expression of the indicated cell surface displayed DNA binding protein, washed, incubated with ssDNA (EC98a) or dsDNA (EC98b_chol), washed and analysed by flow cytometry (Attune NxT, 10,000 cells/sample). Histograms show cell signal in the BL1-H channel, while annotations (%) indicate the proportion of cells considered positive (that score over the indicated gate threshold value) for DNA binding. Overlapping traces show data from triplicate samples. Error bars indicate standard deviation.

**SF4. Displayed p16.7 binds RNA.** Cells (pLO16A) were incubated with (**A.**) ss/dsDNA (EC98b_chol/EC98a, FAM) or (**B.**) ssRNA (EC184_rev_chol/EC184_ fwd, TYE-665), for 30 min at 30^o^C with shaking, before being washed and analysed by flow cytometry. Histograms show cell signal in the BL1-H channel for FAM and RL1-H for TYE-665, while annotations (%) indicate the proportion of cells considered positive (that score over the indicated gate threshold value) for DNA binding. Samples are representative of multiple experiments.

**SF5. Expression of cell surface polymerase.** Constructs for cell surface polymerase expression consisted of a fusion of the phi29 DNA polymerase to the EspP autotransporter (pP2-EspP, SI Table 1), the rest of the vector identical to pSNAP-EspP. The activity of P2-EspP for DNA binding was analysed by flow cytometry. DH10β and C41 strains were transformed with the construct, induced and mixed with cholesterol-linked ss- (**A**) and ds- (**B**) DNA before being washed and analysed by flow cytometry. **C.** Expression of P2-EspP by strain. DH10β and C41 strains (and a DH10β variant with an *ompT* knockout, a gift from Valérie Pezo and Philippe Marlière) were analysed for their expression of P2-EspP (102 kDa) by SDS-PAGE.

**SF6. Cell surface display of T4 DNA ligase. A.** Ligase display constructs, showing leader sequence (orange), ligase variant (yellow), linker (blue) and membrane protein (grey). **B.** Expression of cell surface ligase by construct and strain. DH10β and C41 strains (and a DH10β variant with an *ompT* knockout) were analysed for their expression of T4-EspP (91 kDa) and LO-T4 (69 kDa) by SDS-PAGE. **C-D.** Activity of T4DL for DNA binding by construct and strain. DH10β, C41 and BW25113 *dsbA* strains were transformed with the construct, induced and mixed with cholesterol-linked ss- (**C**) and ds- (**D**) DNA before being washed and analysed by flow cytometry.

**SF7. Quantification of T4DL activity compared to NEB T4DL.** Dilution series of NEB T4 DNA ligase (in red) and T4DL-displaying cells (LOT4, C41, in black) were constructed and mixed with pre-annealed splint ligation substrates MR9/10-F1/11-54PS (as in Fig. 4b) and NEB T4 DNA ligase buffer (1X). Ligation products (37^o^C, 1h; using only the DNA-containing supernatant for the cell samples) were separated by ureaPAGE, imaged (Typhoon imager) and quantified by densitometry (ImageJ). Ligase activity was calculated as the band intensities of: ligated product / (ligated + not ligated species), normalised to 1 for the undiluted NEB T4 DNA ligase. Data is presented as the mean of duplicate experiments (error bars = standard deviation) along with curve-fitting to a Hill function (dashed lines).

**SF8. Flow cytometric analysis of the steps in ligase selection using fluorophore-linked DNA.** Ligase activity using cell surface-linked substrates. Ligation substrates were as described in Fig 4c. Ligation activity was analysed by flow cytometry for T4DL displaying cells (LOT4, C41) before the addition of DNA (*left*), after DNA1 binding(*centre*), and after DNA2 ligation (*right*). Plots show a sample of 1000 cells for clarity (10,000 collected), and indicate the proportion of cells considered positive for (i) DNA attachment (x axis, ROX label), and (ii) Ligation (y axis, FAM label).

**SF9. Ligase selection method using biotin-linked DNA.** Active and inactive displayed ligases were tested in model selection experiments as follows. Cells were subjected to DNA1 binding (grey background), washing to remove nons-transient interactors, followed by DNA2 ligation (green background). Note that DNA2 here is linked to biotin, not FAM. Cells were washed again, and biotin-labelled cells were captured by binding to streptavidin-coated beads as described in the Methods.
