## Supplementary material for "Towards XNA molecular biology: Bacterial cell display as a robust and versatile platform for the engineering of low affinity ligands and enzymes": SI tables: Supplementary Table Legends_v2.docx

**Table S1. Oligonucleotides for cloning.**

Table S1 details the primers used for DNA amplification for each construct, the vectors used for the fragments and the method used for assembly.

**Table S2. Oligonucleotides for DNA binding, ligase assays and ligase selection**

Table S2 details primers used for model selection PCRs as well as DNA binding and ligase assays on the cell surface. DNA modifications are annotated using the same codes used by IDT (<https://www.idtdna.com/pages/products/custom-dna-rna/oligo-modifications>).

**Table S3. Plasmids.**

**Table S4. Strains.**

**Table S5. Gene sequences.** Novel gene sequences that are not available from other sources are provided. LppOmpA and p16.7 were ordered as gblocks, whereas the INP sequence provided is the relevant fragment from plasmid pUC57-INPnc-H.
