## Supplementary material for "Towards XNA molecular biology: Bacterial cell display as a robust and versatile platform for the engineering of low affinity ligands and enzymes": SI tables: Table_S5_v2.docx

**Table S5. Gene sequences**

**LppOmpA**

CCCAATCTAGAAAGGAGATATACACTTCatgaaagcgaccaaactggtgctgggcgcggtgattctgggcagcaccctgctggcgggctgcagcagcaacgcgaaaattgatcagaacaacaacggcccgacccatgaaaaccagctgggcgcgggcgcgtttggcggctatcaggtgaacccgtatgtgggctttgaaatgggctatgattggctgggccgcatgccgtataaaggcagcgtggaaaacggcgcgtataaagcgcagggcgtgcagctgaccgcgaaactgggctatccgattaccgatgatctggatatttatacccgcctgggcggcatggtgtggcgcgcggataccaaaagcaacgtgtatggcaaaaaccatgataccggcgtgagcccggtgtttgcgggcggcgtggaatatgcgattaccccggaaattgcgaccggatgatgagaccttt

**p16.7**

TTTAACTTTAAGAAGGAGATATACCATGGAAGCTATTTTAATGATCGGTGTACTTGCATTGTGCGTTATATTCCTTCTATCAGGACGAAACAACAAAAAGAAACAGGAAGCAAGGGAGCTAGAAGATTATCTTGAAGACCTCAACAAACGAGTTGTTCAACGAACACAAATACTCAGCGAGCTTAACGAAGTTATCTCAAACAGAAGCATTGACAAAACAGTCAACCTGTCAGCTTGTGAAGTCGCCGTGCTTGATCTGTATGAGCAGTCAAATATCCGCATTCCTAGTGACATCATCGAAGATTTGGTTAATCAACGTTTACAAAGTGAACAGGAAGTGTTAAACTATATAGAGACACAGCGGACATACTGGAAATTGGAGAATCAGAAAAAACTATATCGGGGGTCATTGAAATGAGGCCGCACTCGAGCACCACCACCAC

**INP**

ATGACTCTCGACAAGGCGTTGGTGCTGCGTACCTGTGCAAATAACATGGCCGATCACTGCGGCCTTATATGGCCCGCGTCCGGCACGGTGGAATCCAGATACTGGCAGTCAACCAGGCGGCATGAGAATGGTCTGGTCGGTTTACTGTGGGGCGCTGGAACCAGCGCTTTTCTAAGCGTGCAcGCCGATGCTCGATGGATTGTCTGTGAAGTTGCCGTTGCAGACATCATCAGTCTGGAgGAGCCGGGAATGGTCAAGTTTCCGCGGGCCGAGGTGGTTCATGTCGGCGACAGGATCAGCGCGTCACACTTCATTTCGGCACGTCAGGCCGACCCTGCGTCAACGTCAACGTCAACGTCAACGTCAACGTTAACGCCAATGCCTACGGCCATACCCACGCCCATGCCTGCGGTAGCAAGTGTCACGTTACCGGTGGCCGAACAGGCCCGTCATGAAGTGTTCGATGTCGCGTCGGTCAGCGCGGCTGCCGCCCCAGTAAACACCCTGCCGGTGACGACGCCGCAGAATTTGCAGACCGCCACTTACGGCAGCACGTTGAGTGGCGACAATCACAGTCGTCTGATTGCCGGTTATGGCAGTAACGAaACCGCTGGCAACCACAGTGATCTAATTGGATCTGGCGGGCATGACTGCACCCTGATGGCGGGAGAtCAAAGCAGATTGACCGCTGGTAAGAACAGTGTCTTGACGGCAGGCGCTCGTAGCAAACTTATTGGCAGTGAAGGCTCGACGCTCTCGGCTGGAGAAGACTCCACACTAATTTTCAGACTCTGGGACGGGAAGAGGTACAGGCAACTGGTCGCCAGAACGGGTGAGAACGGTGTTGAGGCCGACATACCGTATTACGTGAACGAAGATGACGATATTGTCGATAAACCCGACGAGGACGATGACTGGATAGAGGTAAAGcccggatctgcagaagctgcagctaaAGAAgcagctgcaaaggaagctgcagctaaAGAAgcagctgcaaaGgccatggcAGAAGCGGTCTGA
